# Engineered autocrine signaling eliminates muscle cell FGF2 requirements for cultured meat production

**DOI:** 10.1101/2023.04.17.537163

**Authors:** Andrew J. Stout, Xiaoli Zhang, Sophia M. Letcher, Miriam L. Rittenberg, Michelle Shub, Kristin M. Chai, Maya Kaul, David L. Kaplan

**Affiliations:** Biomedical Engineering Department, Tissue Engineering Resource Center, Tufts University, Medford, MA, USA; Biological Engineering Department, Massachusetts Institute of Technology, Cambridge, MA, USA

**Keywords:** Cultured meat, Cultivated meat, Cellular agriculture, Cell line engineering, Serum-free media, Satellite cells, Fibroblast growth factor, Autocrine signaling

## Abstract

Cultured meat is a promising technology that faces substantial cost barriers which are currently driven largely by the price of media components. Growth factors such as fibroblast growth factor 2 (FGF2) drive the cost of serum-free media for relevant cells including muscle satellite cells. Here, we engineered immortalized bovine satellite cells (iBSCs) for inducible expression of FGF2 and/or mutated Ras^G12V^ in order to overcome media growth factor requirements through autocrine signaling. Engineered cells were able to proliferate over multiple passages in FGF2-free medium, thereby eliminating the need for this costly component. Additionally, cells maintained their myogenicity, albeit with reduced differentiation capacity. Ultimately, this offers a proof-of-principle for lower-cost cultured meat production through cell line engineering.

## Introduction

Cell-cultured meat (also known as cultivated meat or cell-based meat) has the potential to improve the environmental, nutritional, and ethical impacts of our food systems. By decoupling meat from animal husbandry, cultured meat can maintain high control over products and processes to enhance health, efficiency, safety, and sustainability^1–3^. This is particularly true in the context of beef, which presents the largest environmental footprint of industrialized meat production^1^. However, substantial challenges remain before the benefits of cultured meat can be realized, including high costs which inhibit market entry. Currently, these costs are driven by the culture media, which traditionally relies on expensive components such as fetal bovine serum (FBS) or cocktails of recombinant proteins.

Recently, a serum-free medium for bovine satellite cells (BSCs) called Beefy-9 was developed, where costs were driven by recombinant albumin and recombinant fibroblast growth factor 2 (FGF2)^4^. Subsequently, albumin was replaced with low-cost rapeseed (canola) protein isolates to generate the substantially lower-cost medium Beefy-R medium^5^. The dominant cost contributor in Beefy-R is recombinant FGF2 (rFGF), at >60% of the total cost. Therefore, lowering the cost-burden of FGF2 is a clear priority for achieving more affordable culture media.

FGF2 is one of 22 FGFs involved in cell and tissue development and regeneration. This growth factors acts by binding to cell surface FGF receptors (primarily FGFR1) and activating mitogenic pathways. These include the PI3k/Akt pathway (Phosphoinositide 3-kinase / protein kinase B), the Ras/Raf/MEK/ERK pathway (rat sarcoma virus protein / rapidly accelerated fibrosarcoma protein / mitogen-activated protein kinase / extracellular-signal-regulated kinase), and JNK (Jun N-terminal kinase)^6^. In muscle, the activation of these and other pathways maintain growth and inhibit cell fusion^7^. Without FGF2, cell growth decays, and an aberrant phenotype is observed^4^. Thus, exogenous recombinant FGF2 (rFGF) is considered a necessary growth media component for the serum-free culture of BSCs^4,8^.

However, engineering cells for autocrine signaling has the potential to circumvent exogenous growth factor requirements. For instance, overexpression of insulin-like growth-factor 1 (IGF1) and its receptor has been shown to overcome IGF1 requirements in Chinese Hamster Ovary (CHO) cells^9^. Similarly, overexpression of transforming growth factor alpha (TGFα) has been shown to overcome epidermal growth factor (EGF) requirements in colon cells^10^. It has been suggested through preliminary techno-economic assessments of cultured meat that eliminating rFGF from the culture media would reduce production costs at scale by an order of magnitude in some cases^11^. Therefore, the possibility of engineering BSCs for autocrine FGF2 signaling to fully overcome growth factor media requirements is an attractive option.

In this study, the potential for engineered FGF2 autocrine signaling was explored in serum-free culture of immortalized BSCs (iBSCs)^12^. The specific approach was informed by three previously published observations. The first was that MM14 mouse myoblasts naturally express some FGF2, but are still dependent on exogenous FGF2 for proliferation^7,13^. The second was that overexpression of additional FGF2 in MM14s can partially attenuate the dependency on exogenous FGF2, but cannot fully recover growth^14^. Finally, it was observed that overexpression of a constitutively active Ras mutant (Ras^G12V^) can amplify the effects of endogenously expressed FGF2 in MM14 cells to recover the proliferative effects of a low level of exogenous FGF2 *in vitro*, but is not itself sufficient to induce comparable proliferation in FGF2- and serum-free media^15,16^. The authors of this work hypothesized that Ras^G12V^ activates endogenously expressed FGF2, along with offering its own proliferative effect. Together, these observations suggested that engineered overexpression of FGF2 and/or Ras^G12V^ in iBSCs might recover the effects of rFGF, thus allowing for its elimination from the media and substantially reducing costs. In this study, iBSCs^12^ were engineered to express either luciferase (as a control), FGF2, Ras^G12V^, or both FGF2/Ras^G12V^ under a Tet-on promoter (Figure 1).

**Figure 1:**
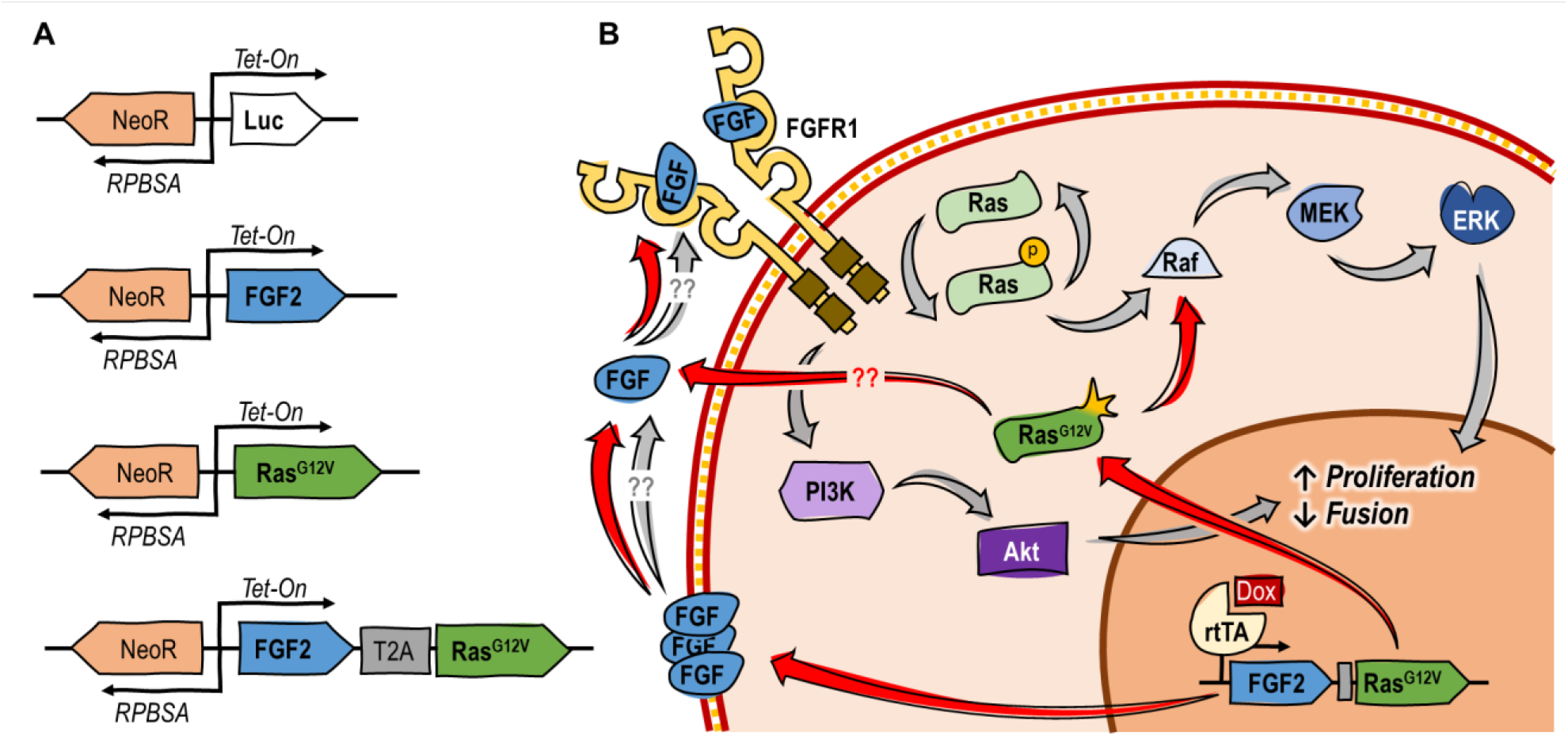
Overview of approach and hypothesized mechanism. (A) Genetic constructs used. (B) Suggested mechanism of action for FGF2/Ras^G12V^, where FGF2 is secreted^17^, complexes with FGFR1, and activates pathways to promote proliferation and inhibit fusion. At the same time, Ras^G12V^ constitutively activates the Ras/Raf/Mek/ERK pathway, and potentially further activates the expressed FGF2^16^.

## Results

### Engineered FGF2 autocrine signaling recovers cell growth in the absence of exogenous FGF

Growth studies in serum-free media with or without exogenous FGF2 (rFGF) showed that control cells were unable to proliferate without rFGF, but cells engineered with FGF2, Ras^G12V^, and FGF2/Ras^G12V^ grew continuously under the same conditions (Figure 2A & S1). Here, the combined expression of both FGF2 and Ras^G12V^ offered improved growth compared with individually expressed genes, and was able to fully recover the proliferation of control cells with 40 ng/mL rFGF. This was the case both when cells were cultured in Beefy-9 without rFGF, and in Beefy-R without rFGF. Over ∼3 weeks, average doubling times in Beefy-9 for controls with rFGF or FGF2, Ras^G12V^, and FGF2/Ras^G12V^ cells without rFGF were 55, 82, 68, and 60 hours, respectively (Figure 2B). This is substantially slower than previously reported growth rates for non-engineered iBSCs grown in serum-containing media (∼15 hrs)^12^, which suggests that future work should focus on adapting iBSCs for serum-free culture or optimizing media and cells to improve growth. During this study, tuning doxycycline concentrations to optimize growth rates was attempted, but offered no improvements (data not shown). Cells engineered for autocrine signaling were also shown to have similar growth whether or not rFGF was present in the media (Figure S1).

**Figure 2:**
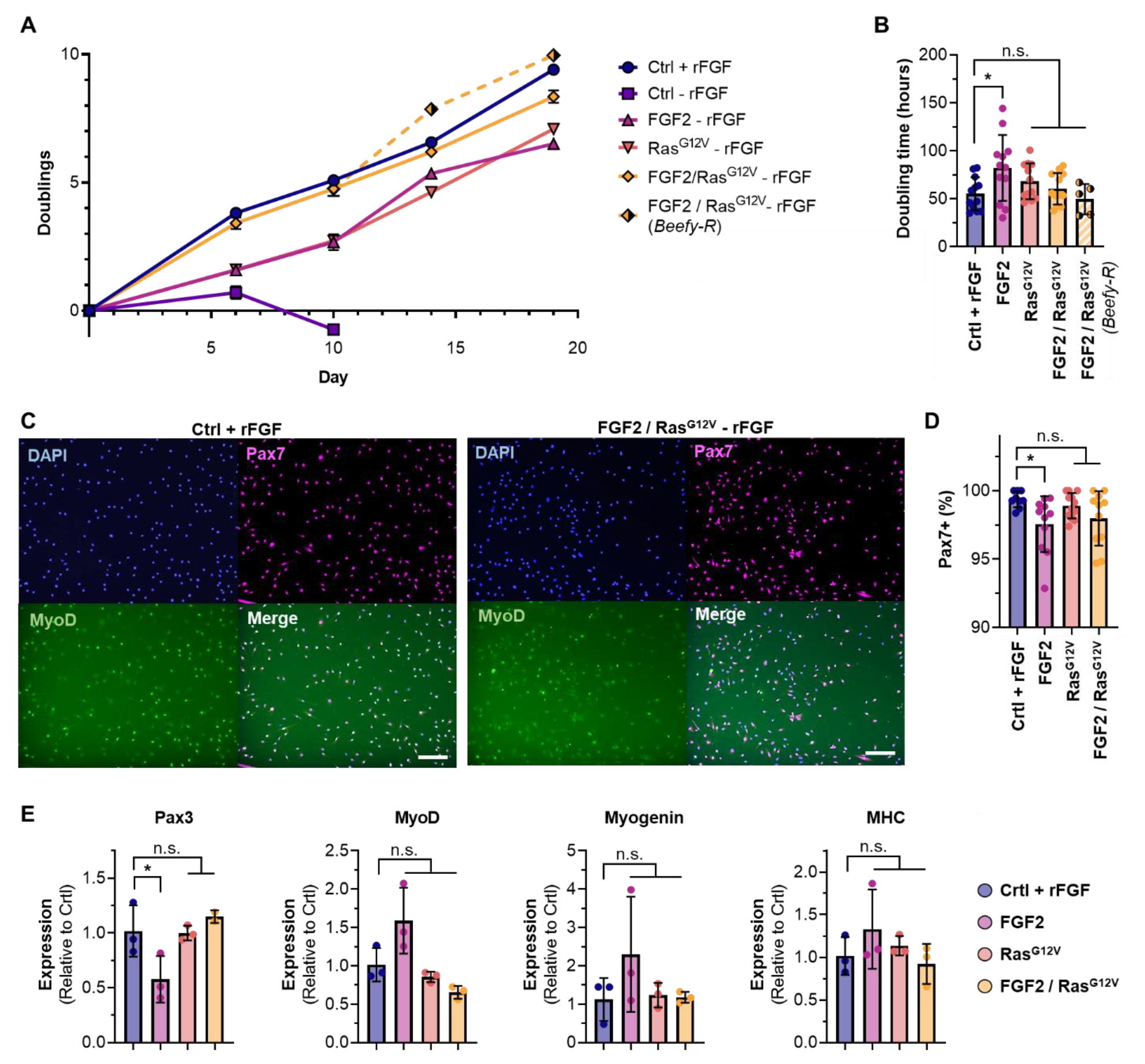
Cell proliferation analysis. (A) Control cells did not grow in the absence of rFGF, while cells expressing FGF2, Ras^G12V^, or FGF2/Ras^G12V^ continuously proliferated under the same conditions. FGF2/Ras^G12V^ cells offered equivalent growth to control cells with rFGF in both Beefy-9 and Beefy-R. n=3. (B) Average doubling times were not significantly different between cell types and controls, except in the case of cells expressing only FGF2. n=12, except in Beefy-R, where n=5; significance is indicated for p <0.05 (*). (C) Immunostaining of controls and FGF2/Ras^G12V^ cells (others in Figure S2). Cells were stained for nuclei (DAPI), Pax7, and MyoD. (D) Analysis of Pax7 staining revealed >97.5% Pax7-positive nuclei in all cell types, with no significant difference between controls and Ras^G12V^ or FGF2/Ras^G12V^ cells. n=9-12, depending on the presence of visual anomalies in images; significance is indicated for p <0.05 (*). (E) Gene expression in proliferating cells relative to controls. Results showed no significant differences except in FGF2 cells, which showed reduced *Pax3* expression. n=3, except in *Pax3* expression of FGF2/Ras^G12V^ cells, where n=2; significance is indicated for p <0.05 (*).

During proliferation, cell phenotypes were assessed via immunostaining for the satellite cell marker Paired Box 7 (Pax7), the early myogenic activation marker myoblast determination protein 1 (MyoD), the early differentiation marker Myogenin, and the terminal differentiation marker Myosin Heavy Chain (MHC), as well as quantitative PCR for *Pax3, MyoD, Myogenin* and *MHC* (Figure 2C-E & Figure S2). Staining revealed high levels of expression of Pax7 and MyoD in engineered cells. Notably, over 97.5% of nuclei in all cell types stained positive for Pax7, and there was no significant difference in Pax7 prevalence between control cells and those expressing Ras^G12V^ or FGF2/Ras^G12V^. These results indicate that engineered iBSCs maintain their satellite cell phenotype. Quantitative PCR revealed that only cells expressing FGF2 alone showed a significant difference (reduced *Pax3*) compared to controls with rFGF. This confirms Pax7 staining results and suggests that cells expressing Ras^G12V^ and FGF2/Ras^G12V^ offer the highest phenotypic similarity to controls.

### Engineered FGF2 autocrine signaling reduces, but does not eliminate, differentiation capacity

Following proliferation, cells were differentiated for two days in a serum-free differentiation medium^18^ without doxycycline and analyzed. Immunostaining for Myogenin and MHC revealed that all cell types maintained the ability to fuse into myotubes, albeit to a reduced level autocrine signaling cells (Figures 3A, 3B, & S3). Gene expression analysis revealed that these Ras^G12V^ and FGF2/Ras^G12V^ cells showed higher expression of *Pax3* than controls, and lower expression of later-stage markers MyoD and Myogenin. Together, these results suggest that autocrine signaling results in a less differentiated phenotype, particularly for cells expressing Ras^G12V^ and FGF2/Ras^G12V^ (Figure 2C). It is possible that additional time could overcome this discrepancy, as cells may still experience lingering signaling two days after the removal of doxycycline. Future work should include optimizing differentiation, as well as proliferation.

**Figure 3:**
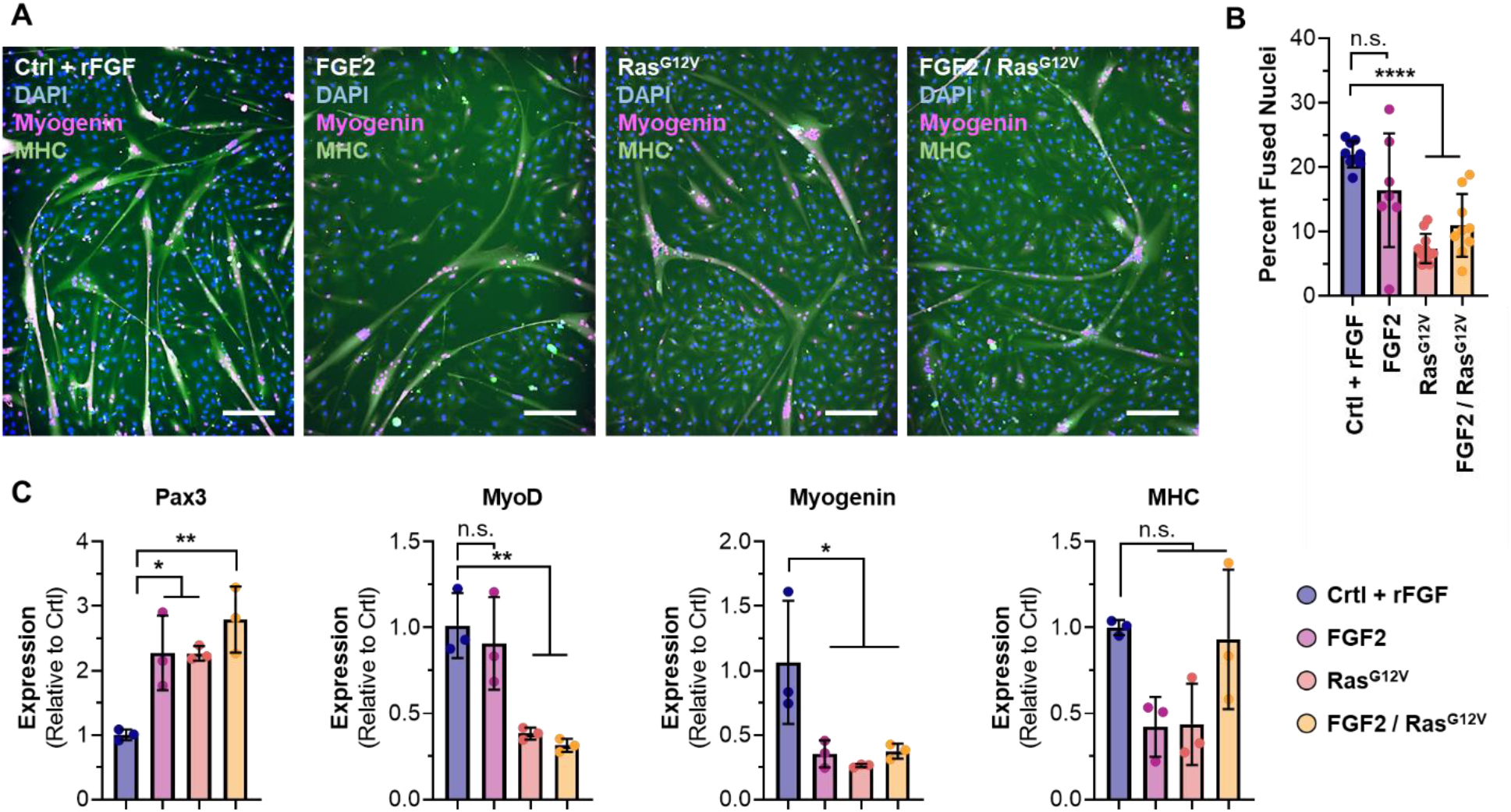
Two-day differentiation of engineered cells. (A) Immunostaining for DAPI, Myogenin and MHC revealed that all cells could fuse into myotubes, but that differentiation capacity was reduced in cells engineered for autocrine signaling. (B) Image analysis revealed that fusion indices of cells engineered for Ras^G12V^ and FGF2/Ras^G12V^ were significantly reduced compared with controls. n=7-11, depending on the presence of visual anomalies in images; significance is indicated for p <0.0001 (****). (C) Gene expression in differentiated cells, relative to controls. Results showed an increase in *Pax3* for autocrine signaling cells, and a reduction in later phenotypic markers. n=3; significance is indicated for p <0.05 (*) and p<0.01 (**).

## Discussion

Lowering the cost of serum-free cell culture media is of paramount importance for cultured meat. Here, we demonstrate a proof-of-principle for reducing costs through engineered autocrine FGF2 signaling in order to overcome costly growth factor requirements (Figure S4). Engineered cells are capable of consistent proliferation in the absence of exogenous rFGF and maintain their satellite cell phenotype. However, reduced growth rates and differentiation capacities for all engineered cells suggest that further optimization of the expression system is required.

Along with optimizing cell lines and expression systems, media composition should be adjusted further in order to maximize process efficiency. For instance, serum-free media other than Beefy-9 and Beefy-R have recently been described for bovine satellite cells, and these media could present promising options for testing and optimizing cell-media systems^8,19,20^. Further optimization should include both basal media components and functional proteins, and can leverage several experimental approaches simultaneously. For instance, computational approaches can increase experimental efficiency to quickly test a large media composition-space^21^. At the same time, metabolomic and proteomic data can be used understand cell behavior in various media, thereby informing directed iterative media formulation efforts^22,23^.

Additionally, further cell line engineering can be used to tune media composition and cell growth and differentiation. For instance, the IGF1 autocrine signaling system discussed previously in CHO cells could potentially overcome insulin requirements in BSCs^9^. Similarly, transferrin overexpression (demonstrated in the same CHO cells) could overcome transferrin requirements in Beefy-9 and Beefy-R. Indeed, by combining these approaches, it is possible that a fully recombinant protein-free culture media could be developed for BSCs and iBSCs, which would dramatically reduce costs^24^. At the same time, it is possible that engineered forms of these various growth factors could amplify their efficacy when expressed endogenously, such as thermally stable and hyper-potent forms of FGF2 or IGF1^25,26^. Lastly, genetic engineering approaches to tune basal media requirements could be explored, such as metabolic engineering of core nitrogen or carbon pathways, or even engineering cells for essential amino acid biosynthesis^27–30^. In each case, tradeoffs between cost-savings and cell culture efficiency will need to be assessed; however, the wide range of possibilities present a valuable toolbox for addressing challenges that could arise during the development, scale-up, and proliferation of cultured meat technologies.

Along with optimization for growth, differentiation, and cost various steps should be taken to render this approach suitable for food. For instance, food-safe induction systems (such as abscisic acid) could be explored to replace doxycycline^31,32^. Similarly, considerations around genetic modifications should be addressed from a regulatory and consumer standpoint. Indeed, there is still substantial consumer concern towards genetically modified foods (GMOs), which in the case of engineered cell lines might exacerbate ambivalent attitudes towards cultured meat^33^. Here, the use of a mutated Ras^G12V^ oncogene, and evidence implicating FGF2 expression with glioma transformation may be important to consider^34,35^. While the Food and Agriculture Organization (FAO) and World Health Organization (WHO) recently found no credible path for cells in cultured meat surviving in the human body or posing tumorigenic risks, consumer perception may still be affected^36^.

Consumer attitudes and regulation of genetically modified cultured meat products will likely face the highest barriers in Europe, which severely restricts the application of genetic engineering for food^37^. This is in contrast to countries such as the USA, which present a more favorable regulatory landscape towards GMOs. Indeed, the Food and Drug Administration (FDA) recently provided a letter of “no further questions” to cultured meat produced with cells that were engineered through cisgenic overexpression of native genes^39^. It is important, here, to note the nuance around how different approaches to genetic engineering are regulated. For instance, the US differentiates transgenic gene insertions from cisgenic insertions, deletions, or base pair changes, which are considered “gene edited” and face fewer regulatory hurdles^38^. As both FGF2 upregulation and Ras mutation are possible using gene editing strategies, such as targeted upregulation or mutation through CRISPR/Cas9, it might be favorable to utilize these approaches when engineering autocrine signaling into practice.

Ultimately, while further development, optimization, and validation of the approach is required, this work provides a proof-of-principle for engineering cells to overcome expensive growth factor requirements. Together with albumin alternatives, this approach can lower the cost of cell expansion, thereby advancing the field towards realizing its potential positive impacts on our food system.

## Acknowledgements

We thank the New Harvest Fellowship Program, the U.S. Department of Agriculture (2021-69012-35978), and the National Institutes of Health (P41EB002520) for supporting this work.

## Author contributions

**AJS**: Conceptualization, Methodology, Investigation, Formal Analysis, Visualization, Writing-original draft preparation, Writing-reviewing and editing. **XZ**: Investigation, Formal Analysis. **SML**: Investigation, Formal Analysis. **MRL**: Investigation. **MS**: Investigation. **KMC**: Investigation. **MK**: Investigation. **DLK:** Conceptualization, Resources, Writing-reviewing and editing, Supervision, Funding acquisition.

## Methods

### Immortalized bovine satellite cell culture and routine maintenance

Immortalized BSCs (iBSCs) used in this study were engineered and characterized previously^12^. For routine cell maintenance, cells were cultured in 37°C with 5% CO_2_ on tissue-culture plastic coated with 0.25 ug/cm^2^ iMatrix recombinant laminin-511 (Iwai North America #N892021, San Carlos, CA, USA). BSC growth media (BSC-GM) was used, comprised of DMEM+Glutamax (ThermoFisher #10566024, Waltham, MA, USA), 20% fetal bovine serum (FBS; ThermoFisher #26140079), 1 ng/mL human FGF2 (ThermoFisher #68-8785-63), 1% antibiotic-antimycotic (ThermoFisher #1540062), and 2.5 μg/mL puromycin (ThermoFisher #A1113803). For routine culture, cells were expanded to a maximum of 70% confluence, harvested using 0.25% trypsin-EDTA (ThermoFisher #25200056), counted using an NC-200 automated cell counter (Chemometec, Allerod, Denmark) and either seeded at a density of 2,000-5,000 cells/cm^2^ on new tissue culture plastic with growth medium and laminin, or else frozen in FBS with 10% Dimethyl sulfoxide (DMSO, Sigma #D2650, St. Louis, MO, USA) and stored in liquid nitrogen.

### Cloning and transfections

The amino acid sequence for bovine *FGF2* was obtained from UniProt (accession number P03969). The gene sequence for this protein was optimized for expression in *Bos taurus* using codon optimization software (IDT, Coralville, IA), and a self-cleaving 2A peptide sequence was added to the end to facilitate bi-cistronic expression. This fragment was ordered through ThermoFisher’s GeneArt gene synthesis service (Table S1). DNA for Ras^G12V^ was obtained from the mEGFP-HRas G12V plasmid, which was a gift from Karel Svoboda (Addgene #18666)^40^. Finally, DNA for neomycin resistance was obtained from the pCDNA3.1 vector (ThermoFisher #V79020).

Genes were inserted into a *Sleeping Beauty* plasmid pSBtet-Pur, which was a gift from Eric Kowarz (Addgene #60507)^41^. Specifically, four new plasmids were generated: 1) pSBtet-Luciferase-NeoR, in which the puromycin resistance of pSBtet-Pur had been replaced with neomycin resistance, 2) pSBtet-FGF-NeoR, in which the luciferase of pSBtet-Luciferase-NeoR had been replaced with FGF2 (minus the 2A peptide), 3) pSBtet-Ras-NeoR, in which the luciferase of pSBtet-Luciferase-NeoR had been replaced with Ras^G12V^, and 4) pSBtet-FGF/Ras-NeoR, in which the Luciferase of pSBtet-Luciferase-NeoR had been replaced by FGF (plus the 2A peptide) followed by Ras^G12V^. In these plasmids, genes of interest (e.g., FGF and/or Ras) were controlled by a Tet-on promoter system, while the Neomycin resistance was promoted by a constitutive RPBSA promoter which ran counter to the Tet-on promoter. Ampicillin resistance and origin of replication were present for standard plasmid maintenance. All cloning was performed via gibson assemblies. This involved amplification with Q5 high-fidelity DNA polymerase (NEB #M0494S, Ipswich, MA, USA), DpnI digestion (NEB #R0176S), PCR cleanup (NEB #T1030S), assembly with the NEBuilder HiFi kit (NEB #E2621S), and heat-shock transformation into competent *E. coli* (NEB #C3019H), according to the suppliers’ instructions. Plasmids were purified with the GeneJet miniprep (ThermoFisher #K0503) and verified via Sanger sequencing (Genewiz, Cambridge, MA, USA). The transposase plasmid which was used for transposon integration was pCMV(CAT)T7-SB100, and was a gift from Zsuzsanna Izsvak (Addgene #34879)^42^.

For transfecting iBSCs, cells were thawed and seeded at 25,000 cells/cm^2^ in 6-well plates with BSC-GM (minus puromycin) and iMatrix laminin-511. The next day, cells were transfected with Lipofectamine 3000 (ThermoFisher #L3000015) according to the manufacturer’s protocol. Briefly, 2.5 μg of each plasmid containing a transposable element was combined with 0.25 μg of the pCMV(CAT)T7-SB100 plasmid, and added to 250 uL of Opti-MEM medium (ThermoFisher #31985088) containing 5 uL of p3000 and 7.5 uL of Lipofectamine 3000. The DNA solution was incubated for 15 minutes at room temperature, during which time cells were rinsed 1X with DPBS and fed 2 mL of Opti-MEM. Next, the DNA solutions were added to the cells, and cells were incubated for six hours at 37°C before 2 mL of BSC-GM (minus puromycin) was added to wells containing Opti-MEM and lipofectamine mixture. Cells were incubated at 37°C overnight, after which media was replaced with BSC-GM supplemented with 2.5 μg/mL puromycin (ThermoFisher #A1113803) and 250 μg/mL G418 (a neomycin analog) (ThermoFisher #10131035). This joint treatment selected for cells which had integrated the new plasmid DNA (e.g., resistant to G418) and maintained the immortalization transgenes (e.g., resistant to puromycin). Cells were selected for fourteen days before being expanded and frozen down for storage. For routine culture, engineered cells were cultured in the puromycin and G418-supplemented media according to the same protocols mentioned above.

### Growth analysis of engineered cells with or without exogenous FGF2

Multi-passage growth studies were performed to validate the utility of FGF2 and Ras^G12V^ expression in serum-free medium lacking recombinant FGF. Briefly, “B8 + FGF”, “B8 - FGF”, “Beefy-9 + FGF”, and “Beefy-9 – FGF” medium were prepared as previously described^4^. The formulations and sources of components can be found in Table S2.

Growth analysis used methods previously established in the development of Beefy-9^4^. Briefly, engineered iBSCs were seeded into triplicate wells of 6-well plates (Corning #353046, Corning, NY, USA) in BSC-GM supplemented with 2.5 μg/mL puromycin, 250 μg/mL G418, and 0.25 ug/cm^2^ iMatrix laminin-511. After allowing cells to adhere overnight, cells were washed 1x with DPBS and fed either Beefy-9 + FGF or Beefy-9 – FGF supplemented with 1μg/mL doxycycline (Sigma #D9891-5G). Media was replaced every two days. For passaging, at 70% confluency, cells rinsed 1x with DPBS, and dissociated with 500 μL of TrypLE Express (ThermoFisher #12604021) at 37°C for 12 minutes. Plates were tapped to dislodge cells, and cells were collected with an additional 1.5 mL of B8 – FGF. Cells were then counted using an NC-3000 automated cell counter (Chemometec), centrifuged at 300 g for 5 minutes, resuspended in either B8 + FGF or B8 – FGF supplemented with 1μg/mL doxycycline (depending on their previous media treatment), re-counted, and seeded onto new 6-well plates at 2,500 cells/cm^2^ with 1.5 μg/cm^2^ of truncated recombinant human vitronectin (Vtn-N; ThermoFisher #A14700). After allowing cells to adhere overnight, media was replaced with either Beefy-9 + FGF or Beefy-9 – FGF supplemented with 1μg/mL doxycycline, as appropriate. This process was repeated over the course of the experiment. For the penultimate passage, a portion of FGF2/Ras^G12V^ cells harvested from the Beefy-9 – FGF condition was seeded onto to additional wells. These cells were fed Beefy-R – FGF supplemented with 1μg/mL doxycycline, in which albumin was replaced with 0.4 mg/mL of a rapeseed protein isolate prepared as previously described^5^.

### Immunostaining (proliferative cells)

Engineered cells were seeded and cultured as before, passaged into 96 well plates, and cultured in the appropriate media (+ doxycycline and +/-rFGF, as appropriate) for two days. Cells were then fixed for 30 minutes with 4% paraformaldehyde (ThermoFisher #AAJ61899AK), washed in DPBS (3x), and permeabilized for 15 minutes with 0.5% Triton-X (Sigma # T8787) in DPBS. Cells were rinsed in PBS-T (3x) comprised of DPBS containing 0.1% Tween-20 (Sigma #P1379) and blocked for 45 minutes using a blocking buffer of DPBS with 5% goat serum (ThermoFisher #16210064) and 0.05% sodium azide (Sigma #S2002). Primary antibodies for Pax7 (ThermoFisher #PA5-68506; 1:500) and MyoD (ThermoFisher #MA5-12902; 1:100) were added, and cells were incubated at 4°C overnight.

The next day, cells were rinsed in PBS-T (3x), incubated in blocking buffer for 15 minutes at room temperature, and treated with secondary antibodies for one hour at room temperature. Secondary antibodies were anti-rabbit (ThermoFisher #A-11072; 1:500) and anti-mouse (ThermoFisher #A-11001; 1:1,000) for Pax7 and MyoD, respectively. A DAPI nuclear stain (Abcam #ab104139, Cambridge, UK; 1:1,000) was included as well. Following incubation with secondaries, cells were rinsed with DPBS (3x). Imaging was performed with a KEYENCE BZ-X810 fluorescent microscope (Osaka, Japan). For Pax7 quantification, batch images were taken with the 10x objective at random points of culture wells selected by the KEYENCE software. Images were analyzed using ImageJ software. Briefly, for Pax7 quantification, each nucleus in the DAPI channel was established as a discrete region of interest (ROI), and these ROIs were added to the Pax7 channel after thresholding at a consistent value. Percentage of ROIs containing Pax7 signal was recorded. Images with clear bubbles or visual anomalies were discarded.

### Immunostaining (Differentiated cells)

Engineered cells were seeded and cultured as before, passaged into 6-well plates, and cultured in the appropriate media (+ doxycycline and +/-rFGF, as appropriate) until reaching 100% confluency. At confluency, cells were rinsed with DPBS (1x), and media was changed to a previously described serum-free differentiation media containing L15 (Invitrogen #11415064) and Neurobasal (Invitrogen #21103049, Carlsbad, CA, USA) media in a 1:1 ratio, supplemented with 10ng/mL IGF1 (Shenandoah Biotechnology #100-34AF-100UG, Warminster, PA, USA), 100 ng/mL EGF (Shenandoah Biotechnology #100-26-500UG), and 1% antibiotic-antimycotic^18^. No puromycin, G418, or doxycycline were added to the differentiation media. Cells were incubated for 2 days before being fixed, permeabilized, and blocked as described for proliferative cells.

Antibodies were then added as before. Primary antibodies for MHC (Developmental studies hybridoma bank #MF-20, Iowa City, IA, USA; 4 μg/mL) and Myogenin (ThermoFisher #PA5-116750; 1:100) corresponded to the anti-mouse and anti-rabbit secondary antibodies mentioned above. A DAPI nuclear stain was added. Cells were rinsed with DPBS (3x) before imaging, and imaging was performed as for proliferative cells. For fusion index quantification, total nuclei were counted in the DAPI channel, after which a consistent threshold was applied to the MHC to generate selection areas. These selection areas were superimposed over the DAPI channel, and nuclei within the selections were counted. The ratios of selected nuclei to total nuclei were recorded as fusion indices. Images with clear bubbles or visual anomalies were discarded.

### Gene expression analysis

Cells were cultured as before in relevant media, and analyzed either at 70% confluency, or after two-days of differentiation as previously described. For the analysis, an RNEasy Mini Kit (Qiagen #74104, Hilden, Germany) was used to harvest DNA according to the manufacturer’s protocol. Next, iScript cDNA synthesis kits (Bio-Rad # 1708890, Hercules, CA, USA) were used to prepare cDNA with 1,000 ng of RNA for each sample, again according to the manufacturers’ instructions. Finally, qPCR was performed using the TaqMan Fast Universal PCR Master Mix without AmpErase UNG (ThermoFisher #4352042). Primers were: *18S* (ThermoFisher #Hs03003631), *Pax3* (ThermoFisher #Bt04303789), *MyoD* (ThermoFisher #Bt04282788), *Myogenin* (ThermoFisher #Bt03258929), and Myosin Heavy Chain (*MHC*) (ThermoFisher #Bt03273061). Reactions were performed using 2 uL of the prepared cDNA, according to the manufacturer’s instructions. Reactions were run on the Bio-Rad CFX96 Real Time System thermocycler (Hercules, CA, USA), and results were analyzed as 2^-ΔΔct^ normalized to 18S expression and comparing relative expression between autocrine signaling cells and control cells.

### Statistical analysis

GraphPad Prism 9.3.0 software (San Diego, CA, USA) was used to perform all statistical tests. All tests were analyzed via one-way ANOVA with a Tukey’s HSD post-hoc test between autocrine engineered cells (without rFGF) and control cells (with rFGF). Valus are given as means, errors are given as ± standard deviation, and p values <0.05 were treated as statistically significant. All replicates shown are distinct samples.

