## Supplementary Material for "Engineered autocrine signaling eliminates muscle cell FGF2 requirements for cultured meat production"

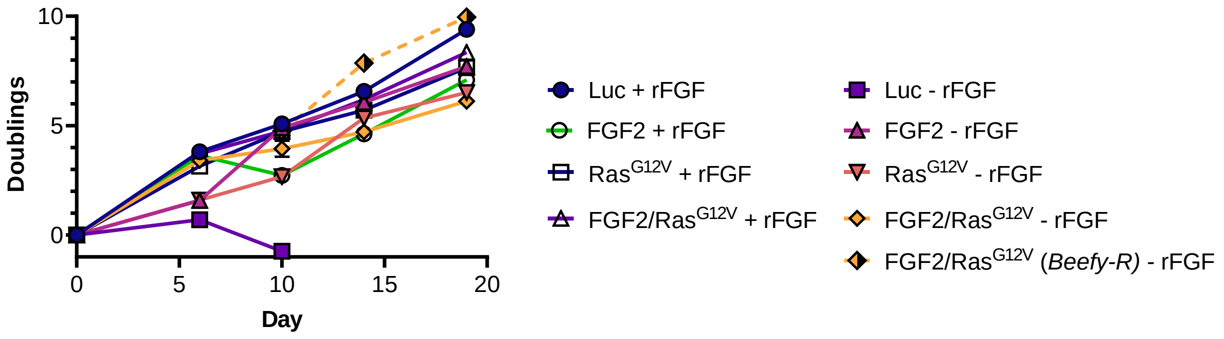


Figure S1: 19-day growth of all engineered cells +/- 40 ng/mL of exogenous recombinant FGF (rFGF). Results showed that growth is maintained for all cells except for luciferase control cells in the absence of exogenous rFGF (Luc – rFGF).

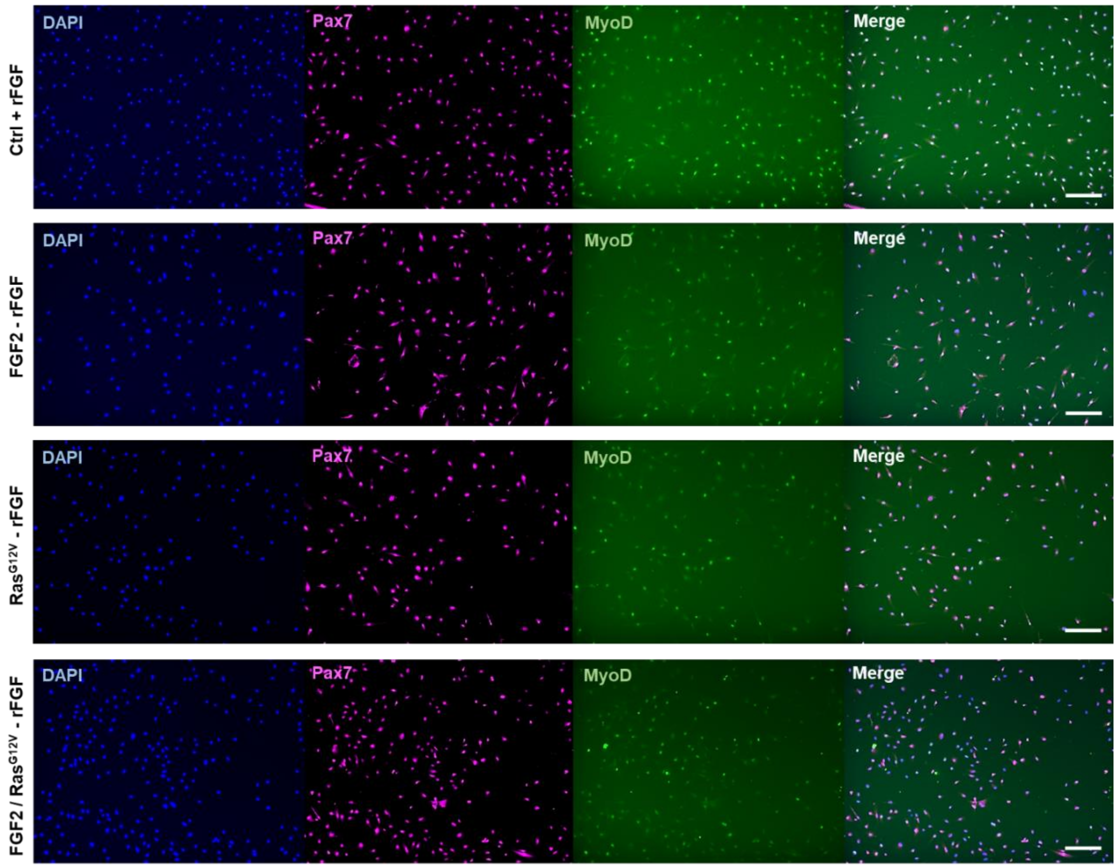


Figure S2: Immunostaining of proliferative cells for nuclei (DAPI), Paired-box 7 (Pax7), and Myoblast determination protein 1 (MyoD). Scale bars 200 µm.

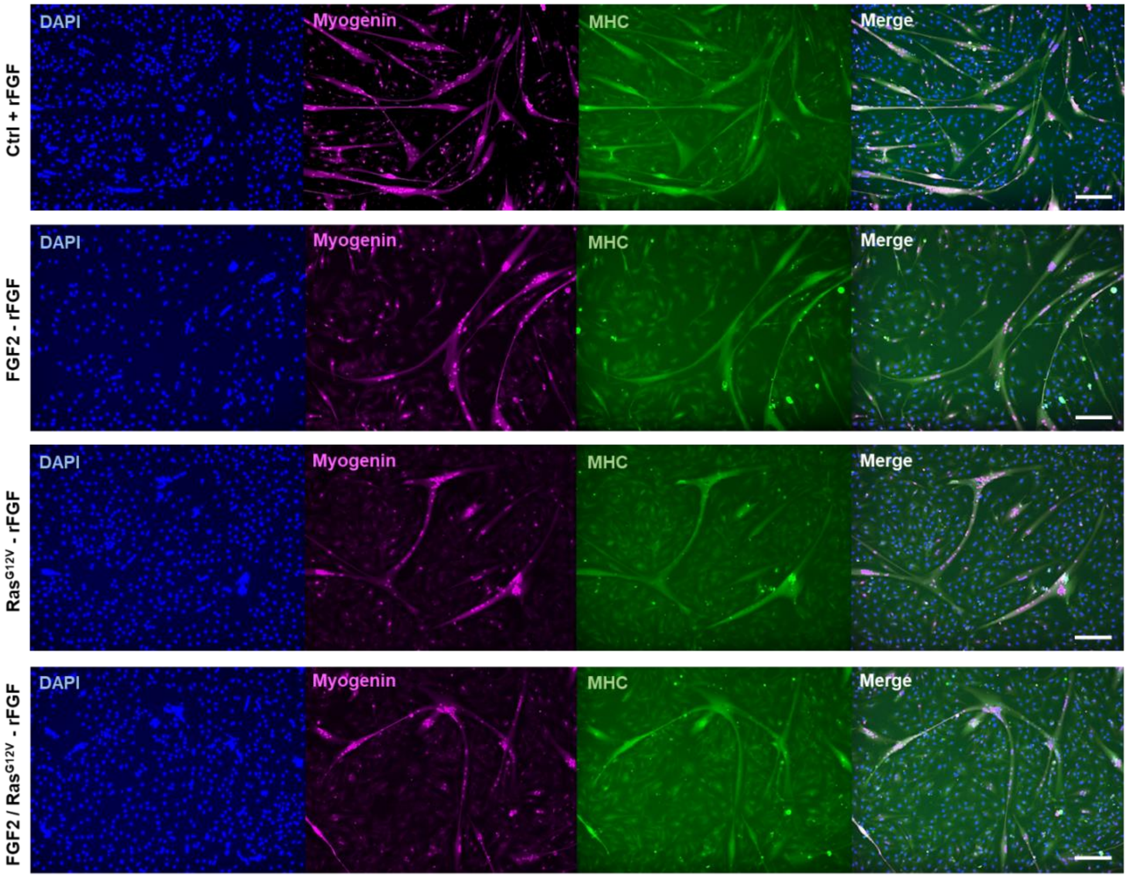


Figure S3: Immunostaining of differentiated cells for nuclei (DAPI), Myogenin, and Myosin Heavy Chain (MHC). Scale bars 200 µm.

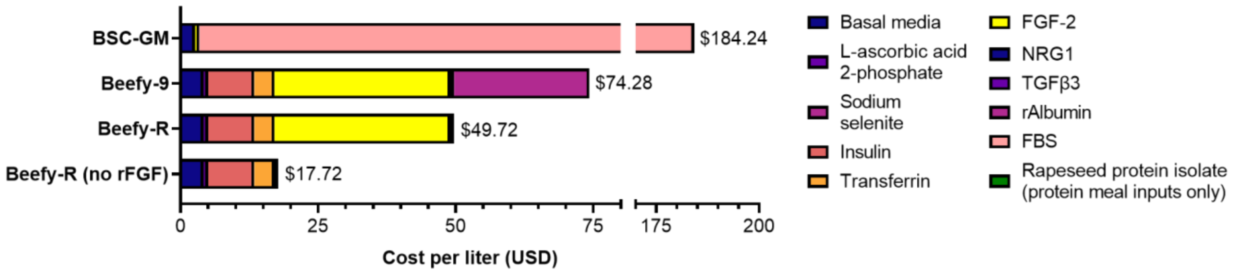


Figure S4: Cost analysis of serum-containing media (BSC-GM), serum-free Beefy-9 media, serum-free Beefy-R media (in which rAlbumin was replaced with rapeseed protein isolate), and Beefy-R without rFGF, which is made possible through autocrine engineering. Media prices based on bulk ordering of components described previously[1,2].

| | |
|---|---|
| FGF2 | ATGGCGGCAGGTTCAATAACTACCCTGCCTGCTCTGCCTGAGGATGGCGGATCTGGTGCTTTTCCACCTGG<br>CCACTTCAAGGACCCCAAGAGGCTGTACTGCAAGAACGGCGGATTCTTCCTGAGGATTCACCCCGACGGA<br>AGAGTGGATGGCGTGCGCGAAAAGAGCGACCCTCACATCAAGCTGCAACTGCAGGCCGAAGAGAGAGGC<br>GTCGTCAGCATCAAAGGCGTGTGCGCCAATAGATACCTGGCCATGAAGGAAGATGGCAGGCTGCTGGCCT<br>CCAAATGCGTGACCGACGAGTGCTTCTTCTTCGAACGCCTGGAAAGCAACAACTACAACACCTACAGAAG<br>CCGCAAGTACTCCTCTTGGTACGTGGCCCTGAAGAGGACCGGCCAGTACAAGCTGGGACCTAAGACAGGC<br>CCTGGACAGAAGGCCATCCTGTTCCTTCCAATGTCCGCCAAAAGTggatctggcgaaggcagaggctctctgctgacatgtggcga<br>cgtggaagagaaccctggacct |
| Ras$^{G12V}$ | ATGACGGAATATAAGCTGGTGGTGGTGGGCGCCGTCGGTGTGGGCAAGAGTGCGCTGACCATCCAGCTGA<br>TCCAGAACCATTTTGTGGACGAATACGACCCCACTATAGAGGATTCCTACCGGAAGCAGGTGGTCATTGA<br>TGGGGAGACGTGCCTGTTGGACATCCTGGATACCGCCGGCCAGGAGGAGTACAGCGCCATGCGGGACCA<br>GTACATGCGCACCGGGGAGGGCTTCCTGTGTGTGTTTGCCATCAACAACACCAAGTCTTTTGAGGACATCC<br>ACCAGTACAGGGAGCAGATCAAACGGGTGAAGGACTCGGATGACGTGCCCATGGTGCTGGTGGGGAACA<br>AGTGTGACCTGGCTGCACGCACTGTGGAATCTCGGCAGGCTCAGGACCTCGCCCGAAGCTACGGCATCCC<br>CTACATCGAGACCTCGGCCAAGACCCGGCAGGGAGTGGAGGATGCCTTCTACACGTTGGTGCGTGAGATC<br>CGGCAGCACAAGCTGCGGAAGCTGAACCCTCCTGATGAGAGTGGCCCCGGCTGCATGAGCTGCAAGTGTG<br>TGCTCTCCTAA |

Table S1: Genes used. Gene sequences used in constructs (upper case), followed by 2A linker sequences used after FGF-2 for bi-cistronic expression with Ras$^{G12V}$ (lower case).

| Component | Concentration | Supplier | Catalog # |
|---|---|---|---|
| **DMEM/F12 basal media** | **N/A** | **ThermoFisher** | **11320033** |
| **2-Phospho-L-ascorbic acid trisodium salt** | **200 µg/mL** | **Sigma** | **49752-10G** |
| **Insulin (human, recombinant)** | **20 µg/mL** | **Sigma** | **91077C-250MG** |
| **Transferrin (human, recombinant)** | **20 µg/mL** | **InVitria** | **777TRF029** |
| **Sodium selenite** | **20 ng/mL** | **Sigma** | **S5261-10G** |
| **Fibroblast growth factor (FGF-2)** | **40 ng/mL** | **PeproTech** | **100-18B** |
| **Neuregulin (NRG1)** | **0.1 ng/mL** | **PeproTech** | **100-03** |
| **Transforming growth factor (TGFβ3)** | **0.1 ng/mL** | **R&D Systems** | **8420-B3-005/CF** |
| **UltraPure Water** | **5.8% (v/v)** | **ThermoFisher** | **10977015** |
| **Antibiotic/Antimycotic** | **1% (v/v)** | **ThermoFisher** | **1540062** |
| **Recombinant albumin** | **0.8 mg/mL** | **Sigma** | **A9731-1G** |

Table S2: Components of the various media used throughout this study. “Beefy-9 + FGF” had all components, “Beefy-9 – FGF” had all components except FGF-2, “B8 + FGF” had all components except recombinant albumin, and “B8 – FGF” had all components except FGF-2 and recombinant albumin.
